# Relative changes in the cochlear summating potentials to paired-clicks predict speech-in-noise perception and subjective hearing acuity

**DOI:** 10.1101/2022.07.31.502232

**Authors:** Jesyin Lai, Gavin M. Bidelman

## Abstract

Objective assays of human cochlear synaptopathy (CS) have been challenging to develop. It’s suspected if relative summating potential (SP) changes are different in CS listeners. In this proof-of-concept study, young, normal-hearing adults were recruited and assigned to a low/high-risk group for having CS based on their audiograms at 9-16 kHz. SPs to paired-clicks with varying inter-click intervals isolated non-refractory receptor components of cochlear activity. Abrupt increases in SPs to paired- vs. single-clicks were observed in high-risk listeners. Critically, overexaggerated SPs predicted speech-in-noise and subjective hearing abilities, suggesting relative SP changes to rapid clicks might help identify putative synaptopathic listeners.

## 1. Introduction

Recent animal studies reveal that intense noise overexposure can lead to cochlear neuronal degeneration, even when hair cells recover and thresholds return to normal^1^. Up to 50 % loss of synapses between inner hair cells and cochlear nerve fibers can occur in noise-exposed or aging ears without prominent hair cell loss or auditory threshold elevation^2,3^. Hence, this cochlear synaptopathy is “hidden” because it is not detected by routine behavioral or electrophysiological measures (e.g., clinical audiogram thresholds)^4^. Moreover, although loss of cochlear synapses can happen immediately after extreme noise exposure, the subsequent degeneration of the cochlear nerve fibers develops over a more protracted time course^1^. Neural degeneration following cochlear synaptopathy is thought to manifest in poor word-recognition scores^3,5–7^, particularly in noise^8–11^, but not pure-tone detection in quiet. This is partly because cochlear neurons with high thresholds and low spontaneous rates are preferentially affected by cochlear synaptopathy^12,13^, though this does not applied to all species^14^.

One common way to diagnose cochlear synaptopathy in noise-exposed or aged animals with normal-hearing thresholds is by using the suprathreshold amplitude of auditory brainstem response (ABR) wave I, which represents the summed activity of cochlear neurons and correlates with loss of cochlear synapses in corresponding frequency regions^1,2,15^. However, using ABR wave I amplitude as a diagnostic tool in humans is challenging^16,17^ and has provided mixed results^18^. The low amplitude response is highly variable and sometimes unmeasurable at the scalp. On the other hand, the summating potential (SP), reflecting aggregated hair cell receptor activity^19^, is observed to be larger in listeners with poor speech-in-noise scores^7^ or with greater acoustic overexposure^8^. SP enhancement was also reported in listeners immediately after recreational acoustic overexposure^20^. Given that the SP reflects hair cell-dendritic integrity rather than the output of the auditory nerve (cf. Wave I), it may be more sensitive to synaptopathic pathology. In normal-hearing human listeners, SP amplitudes to short duration stimuli are found to be reducing as stimulus repetition rate increased^21^. In contrast, as excitatory post-synaptic potentials (EPSPs) from cochlear nerve terminals under the inner hair cells may contribute to SP^8^, the loss of a negative EPSP could enhance SP in cochlear synaptopathy. SP enhancement is also reported in synaptopathic mice with attenuated middle-ear-muscle reflex (MEMR)^22^. However, like ABR wave I, a critical limitation to overcome is the low-amplitude nature of the SP, which hinder the use of absolute measures of the response as a viable diagnostic.

Here, in this proof of concept study, we evaluated whether relative changes in SP to rapid auditory stimuli might serve as a new potential assay of cochlear synpatopathy. We recruited young participants with normal and similar hearing thresholds in the normal audiometric range (250-8000 Hz) but who differed in their extended high-frequency (EHF) thresholds (9-16 kHz). We divided them into low- and high-risk groups based on their average EHFs thresholds of both ears. We then measured SPs elicited by standard single-clicks and paired-clicks^23^ (in separate stimulus conditions). In this paired-click paradigm, two clicks were presented within a short interval (e.g., 0.1-4.0 ms) to boost the generation of SP. As ABR wave I is the sum of action potentials generated by auditory neurons upon activation by a stimulus, there is a refractory period in which neurons are not excitable again in response to another stimulus. Unlike ABR wave I, SP is not limited by refractoriness in response to the first click of a pair. However, SP elicited by the second click of a pair is still regulated by synaptic processes due to neurotransmitter release^24^ and re-uptake^25^. When a click is placed within 1 ms of a preceding click (i.e., within the absolute refractory period of the auditory nerve), short-term adaptation inhibits a (neural) response to the second click, which should minimize the ABR to the second click and isolate activities of (pre-neural) afferent spiral ganglion dendrites^23^. Hence, SP stimulated with double-clicks may provide a more “pure” measurement of receptor and dendritic response integrity. Instead of measuring absolute SP amplitude, we were also interested to test whether *relative* measures of the SP to rapid auditory stimuli, which minimize inter-subject differences, would provide a more sensitive assay of putative cochlear synaptopathy in humans.

## 2. Materials and Methods

### 2.1 Participants

The sample included N = 18 young participants with age ranging 23-33 years (M = 25, SD = 2.8 years; 10 females). All spoke American English and had normal hearing (20 dB HL; 250–8000 Hz) when tested by conventional audiometric standards. Each gave written informed consent in compliance with the University of Memphis IRB.

### 2.2 Auditory Test Battery

We obtained subjective hearing acuity and noise exposure history of each participant by using the Lifetime Exposure to Noise and Solvents – Questionnaire (LENS-Q)^26^, and two additional noise questionnaires [see in Appendix 1 of Liberman et al. (2016)]. Moreover, we measured otoscopy, ipsilateral acoustic reflex thresholds (ARTs), and distortion product otoacoustic emissions (DPOAEs) from participants’ right ears according to standard audiological conventions. The final ART was defined as the averaged threshold obtained across the four elicitor frequencies (0.5, 1, 2 or 4 kHz).

### 2.3 Audiometric thresholds and QuickSIN

We measured pure-tone air-conduction thresholds bilaterally from 250 Hz to 16,000 Hz at octave intervals and also at 9, 10, 11.2, 12.5 and 14 kHz. For 8 kHz and below, ER-3A inserts were used for testing; for extended high frequencies (EHFs) above 8 kHz, circumaural headphones were used (Sennheiser HDA 200) that were specialized for high-frequency audiometry. We divided participants into two groups, low- vs. high-risk, based on their average EHF thresholds. High-risk participants (N = 9, 4 females) had an average EHF threshold of 12.9 ± 8.24 dB hearing level (HL) while low-risk participants (N =9, 6 females) had an average EHF threshold of −1.71 ± 2.91 dB HL.

To assess participants’ speech perception in noise, we used the Quick Speech-in-Noise (QuickSIN) test^27^. Participants heard lists of 6 sentences, each with 5 target keywords spoken by a female talker embedded in four-talker babble noise. Target sentences were presented binaurally at 70 dB sound pressure level (SPL) at signal-to-noise ratios (SNRs) decreasing in 5 dB steps from 25 dB (relatively easy) to 0 dB (relatively difficult). SNR-loss scores reflect the difference between a participant’s SNR-50 (i.e., SNR required for 50 % keyword recall) and the average SNR threshold for normal hearing adults (i.e., 2 dB)^27^. Higher scores indicate poorer SIN performance. Participants’ scores ranged from −4 to 2 dB of SNR-loss (M = −0.03, SD = 1.74). High-risk participants’ mean score was 0.6 ± 1.19 SNR-loss, whereas low-risk participants’ mean score was −0.6 ± 2.02 SNR-loss (i.e., 1 dB better performance). The QuickSIN scores of both groups fell within the normative range for normal-hearing individuals.

### 2.4 Stimuli and electrophysiology

Standard single-clicks with a duration of 0.1 ms and inter-stimulus interval of 125 ms (repetition rate = 8 Hz) were presented monaurally at 80 dB SPL to participants’ right ears. Acoustic stimuli were delivered in alternating polarity via ER-3A earphones. A vertical montage (Fpz to tiptrode) was used to record ABRs. A tiptrode (ear canal) reference was used to enhance the more peripheral (i.e., cochlear) components of the electrocochleogram (ECochG) and better visualize the pre-synaptic SP^8,28^. Inter-electrode impedance was kept below 2 kΩ. Electrical responses were amplified 100,000x, sampled at 10 kHz, and filtered with a 50–3,000 Hz band-pass plus 60 Hz notch filter. Two thousand five hundred artifact-free sweeps were averaged to obtain low-noise ECochGs.

Following standard single-clicks, we also recorded ECochGs from participants using a paired-click paradigm^23^. In this paradigm, paired-click stimuli with 7 different inter-click intervals (ICI) of 4.0, 2.0, 1.0, 0.8, 0.4, 0.2, and 0.1 ms were presented (in separate stimulus sessions and random condition order). Stimulus delivery and recording settings were kept the same as the single-clicks. The time intervals were chosen to be shorter than, encompass and exceed the duration of absolute (1.0-1.2 ms) and relative (4-5 ms) refractory periods of the auditory nerve^29^. This resulted in a total of 8 ECochG/ABR waveforms per listener.

### 2.5 ECochG waveform analysis

All ECochG waveforms were processed and analysed using customized scripts in Python 3.9.7. First, each waveform per participant and condition (epoched from −12 to 12 ms relative to the first click onset) was low-pass filtered at 2500 Hz to remove high-frequency noise. For waveform, we then performed baseline correction by obtaining an average amplitude at −5-0 ms and subtracting this average amplitude from all amplitudes within the epoch window. Next, we defined SP time range as 0.5-1.3 s post click onset as the reported mean latency of SP is 0.97 (± 0.1) ms while wave I mean latency is 1.83 (± 0.1) ms^23^. This SP time range and latency are also consistent with definitions in other recent studies on ECochG/ABR and synaptopothy^8,9^. Subsequently, we fit a logarithmic curve (refer to Fig. 2A) using log*(SP time) and original amplitudes of SP time by performing a polynomial fitting (degree=1) with ‘numpy’ package in Python (Eq. 1):

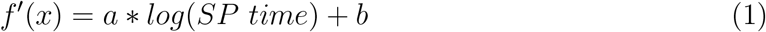

**Fig. 1.**
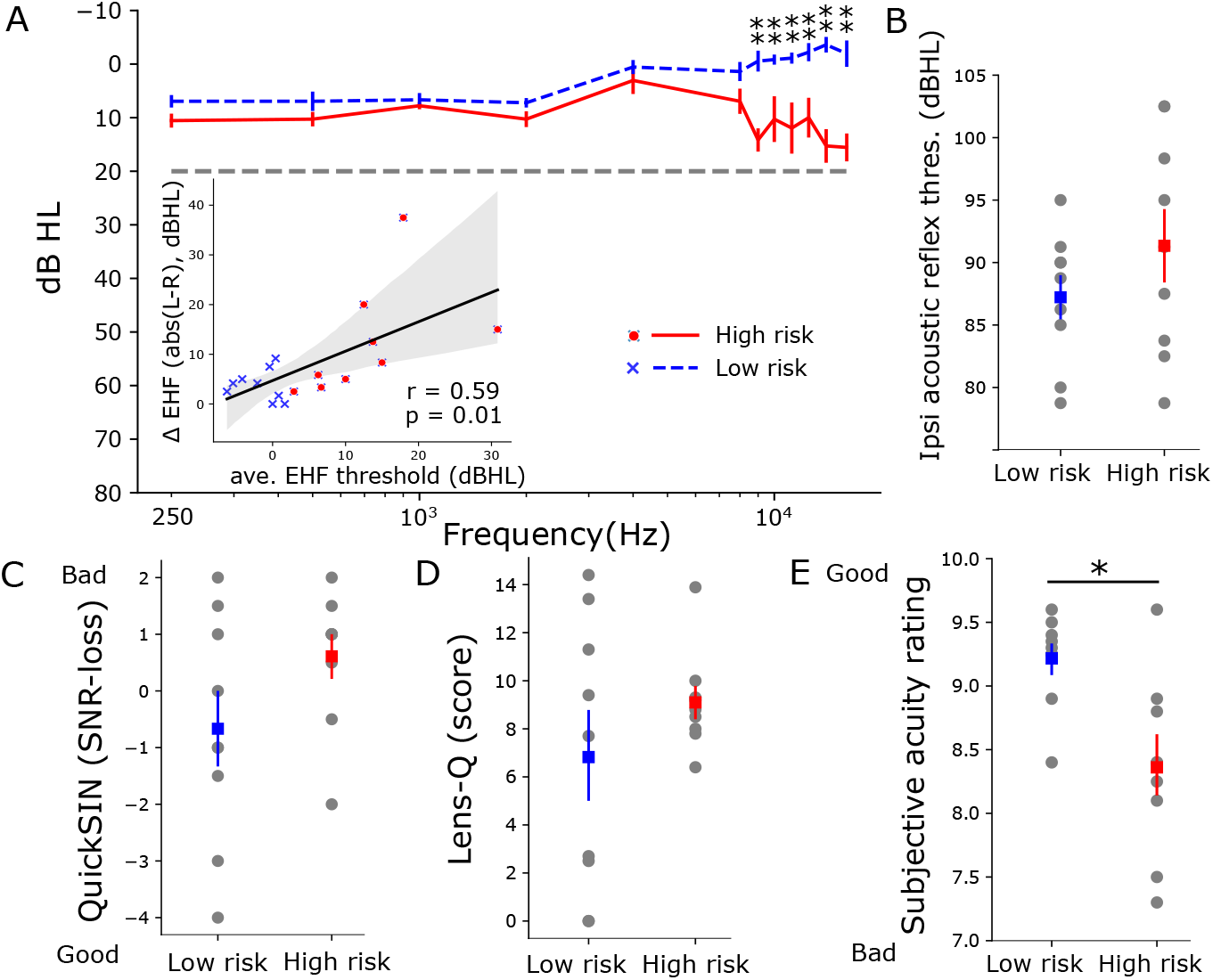
High-risk listeners have poorer and more asymmetric auditory thresholds at extended high-frequencies (>8 kHz) and poorer subjective acuity than low risk listeners. (A) Low- and high-risk groups have similar auditory thresholds in the normal audiometric range (250-8000 Hz) indicative of “hidden” hearing loss. For extended high frequencies (EHFs, 9-16 kHz), thresholds of high-risk participants were significantly poorer (Mann Whitney U test) than low-risk participants. The inset in (A) shows the strong association of average EHF threshold differences between left and right ears (Δ EHF) with average EHF thresholds of both ears. Despite trends, risk groups did not differ in (B) ipsilateral acoustic reflex thresholds, (C) QuickSIN, nor (D) LENS-Q noise exposure scores. (E) Subjective acuity ratings were better in the low- vs. high-risk group (Mann Whitney U test, U-val= 69, p = 0.01). Errorbars= ± s.e.m., *p<0.05 and **<0.005, r = Spearman’s correlation, shaded area indicates 95 % confidence interval of the regression line.

**Fig. 2.**
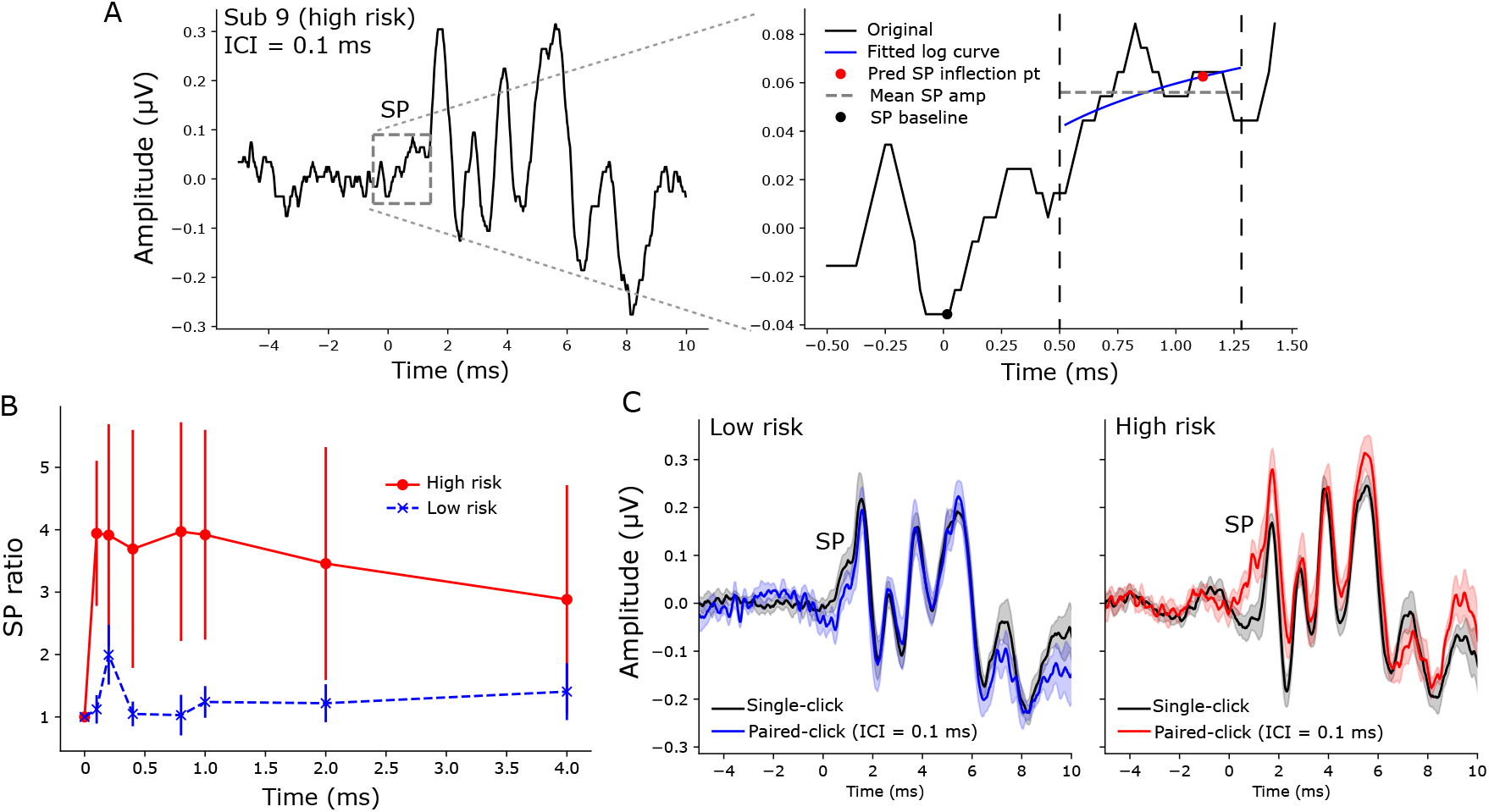
Summating potential (SP) ratio (SP amplitudes to paired-clicks/SP amplitudes to singleclicks) were larger in the high-compared to low-risk group. (A) An example of ECochG/ABR recorded from a representative high-risk subject (left panel; paired-click response, ICI=0.1 ms). The SP is labeled with a gray box and enlarged in the right panel. A logarithmic curve (blue line) was fit from 0.5 to 1.3 ms (the region within two dashed vertical lines) to predict the SP inflection point (red circle) at 85 % of min-max amplitude. SP amplitude was measured as the difference between the predicted SP inflection point and the SP baseline (min. amplitude within −0.5-0.5ms, black circle). In some cases when the fitted curve had a slope of ≤0, SP amplitude was measured as the difference of the mean amplitude within 0.5-1.3 ms (gray dashed line) and the SP baseline. (B) Increased SP amplitudes to paired-clicks was observed in high-risk participants when normalized to the SP amplitude of single-click (within each participant). (C) Grand average ECochGs to single- and paired-clicks (ICI = 0.1 ms) in low- (left) and high-risk (right) groups. Errorbars and shaded area = ± s.e.m.

Where *SP time* is the sampled time vector within 0.5-1.3 ms, *a* is the fitted slope, *b* is the fitted constant and *f*′(*x*) is the predicted amplitudes of SP time. The SP inflection point was estimated at the 85 % the min-max rise of the fitted curve, as:

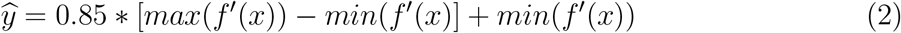

Where *ŷ* is the estimated SP inflection point. Finally, SP amplitudes were defined as the difference between the estimated SP inflection point and SP baseline, which was defined as the lowest amplitude within −0.5 - 0.5 ms (cf.^8^). Meanwhile, if the fitted slope (i.e., a) was ≤0, SP amplitude was defined as the mean of original amplitudes in SP time subtracting SP baseline. In a few instances where SP amplitudes were lower than SP baseline, SP measures were excluded and considered no response (i.e., missing data).To measure relative changes in SP with increasing click ICI, SP amplitudes for each ICI condition were normalized to the SP amplitude of single-clicks per participant to obtain an SP ratio. This differential metric allowed us to assess changes in the SP within each listener, thus avoiding confounds of absolute measures.

### 2.6 Statistical analysis

To compare audiometric thresholds and behavioral measures between low- and high-risk groups, we used non-parametric Mann Whitney U test (‘pingouin’ package in Python). We performed a two-way, mixed-model ANOVA (participants = random factor) using ‘lme4’ package in Rstudio to compare SP ratios across the two main factors (ICI; risk group). Initial diagnostics were performed using residual and Q-Q plots to assess heteroscedasticity and normality of data. Effect sizes are reported as *η_p_*^2^. To assess pairwise linear relations between electrophysiological measures (i.e., normalized SP amplitudes or SP ratios) and behavioral measures (i.e., QuickSIN, LENS-Q, average EHF thresholds, average EHF threshold differences between ears, subjective acuity rating, and ARTs), we used Spearman’s correlations (‘scipy’ package in Python).

## 3. Results

### 3.1 Audiometry, QuickSIN, and self reported hearing acuity

Behavioral measures including normal and EHF thresholds, ARTs, QuickSIN, LENS-Q scores, and subjective acuity ratings are shown for low- vs. high-risk listeners in Fig. 1. Average EHF thresholds of the high-risk group were worse (Mann Whitney U test, U-val= 0, p < 0.05) than the low-risk group although auditory thresholds of both groups fell within the clinically normal range (i.e., better than 20 dB HL). The high-risk group also showed more than 12 dB asymmetry between left and right thresholds for 9-16 kHz. Asymmetry in EHFs, however, was only 3.8 dB in the low-risk group. Asymmetry in EHFs associated strongly with average EHF thresholds (Spearman’s r= 0.59, p = 0.01). Meanwhile, differences in ipsilateral ARTs, QuickSIN, and LENS-Q scores were not significant when compared across the low- and high-risk groups (Fig. 1B-D). Average subjective acuity rating (Fig. 1E) was significantly lower (i.e., poorer hearing sensitivity, U-val=0.69, p = 0.01) in the high-risk group.

### 3.2 Electrocochleography

SP amplitudes, which reflect hair cell receptor potentials^19,30^, evoked by standard single- and paired-clicks (ICI = 0.1 ms) are shown in Fig. 2C. An example of SP quantification using curvefitting (refer to Methods and materials) is shown in Fig. 2A. We normalized SP amplitudes of paired-clicks to SP amplitudes of single-clicks per participant to lessen the impact of the inter-subject differences in head size, electrode contact, etc., which confound absolute measures and ECochG amplitude. SP ratios were generally larger (F_1,16_ = 4.47, p =0.05, *η_p_*^2^ = 0.22) in the high-risk than the low-risk group across ICI conditions (Fig. 2B). Grand average waveforms clearly show the prominent increase in SP amplitude in the high-risk group when evoked by paired-clicks (right panel of Fig. 2C).

### 3.3 Brain-behavior associations

We ran correlations to determine if SP ratios across ICI conditions (0.1-4.0 ms) were related to behavior. Fig. 3A-B show that SP ratios at ICI = 0.1 (Spearman’s r = 0.57, p = 0.03) and 0.2 ms (r = 0.6, p = 0.02) were predictive of QuickSIN scores; poorer speech-in-noise performance was associated with larger (i.e., overexagerated) SP ratios. Furthermore, SP ratios at ICI = 1.0 ms were predictive of subjective acuity ratings (Fig. 3C). We also found that subjective acuity ratings had strong associations with average EHF thresholds and average EHF threshold differences between ears (Fig. 3D-E). These results indicate links between poorer subjective hearing acuity and (i) poorer high-frequency hearing abilities and (ii) more asymmetric high-frequency hearing thresholds. Additional correlation analyses treating average EHF threshold or average inter-aural EHF threshold difference as a continuous variable did not reveal significant associations with SP ratios at various ICIs though trends of positive associations were observed. Similar correlation analyses were also performed on SP ratios at all ICIs and other behavior measures (e.g., LENS-Q, ipsilateral ART, etc.) but not all correlations were significant and only significant correlations were reported over here.

**Fig. 3.**
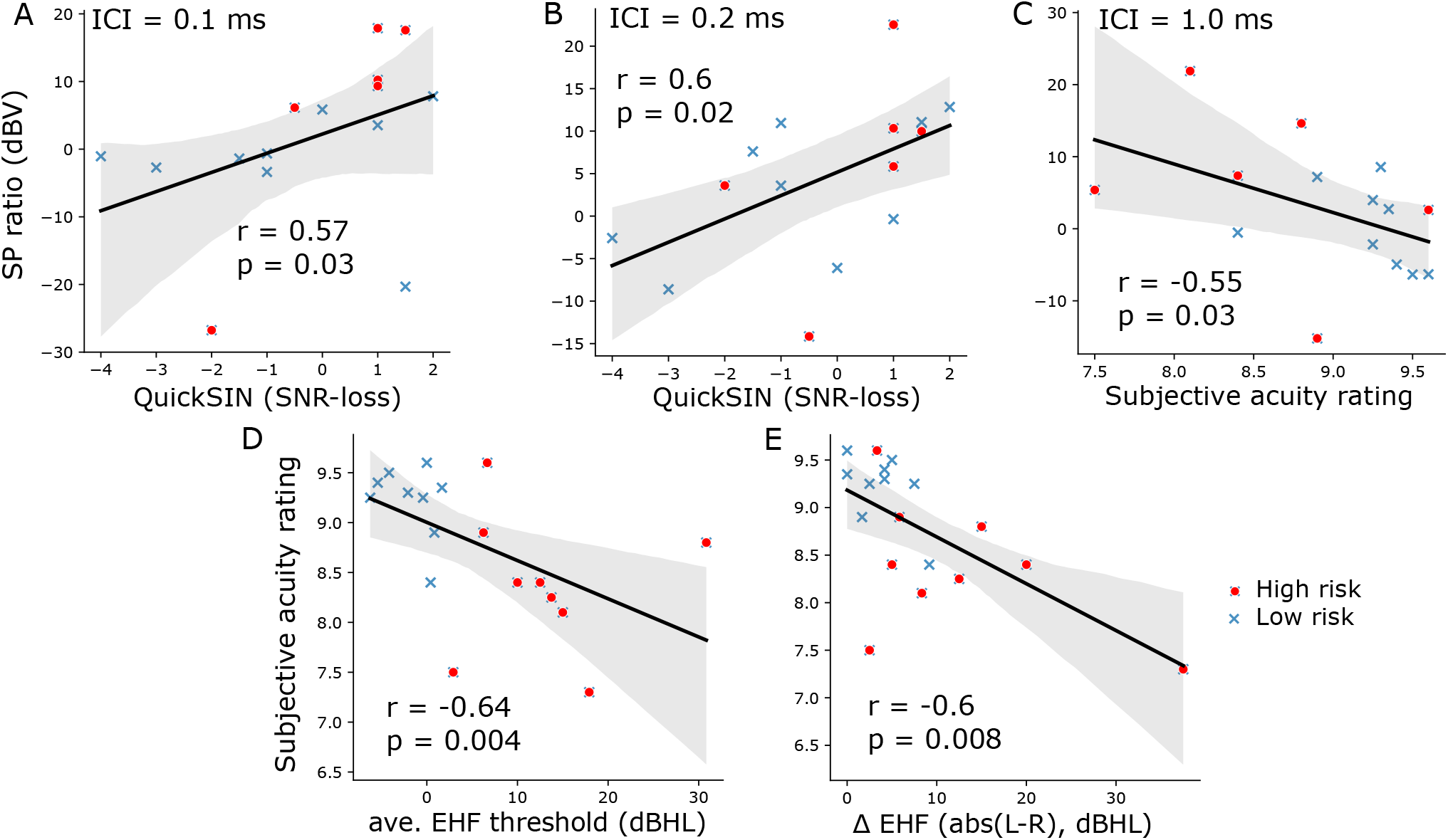
Brain-behavior correlations. (A-B) SP ratios are related to speech-in-noise perception. SP ratios (plotted in dBV scale) of (A) ICI = 0.1 ms and (B) 0.2 ms correlated significantly with QuickSIN SNR-loss. (C) Largrer SP ratios for 1.0 ms predicted poorer subjective acuity rating. (D-E) Extended high-frequency (EHF) hearing abilities are related to subjective hearing acuity. Subjective acuity rating was strongly predictive of (D) average EHF thresholds and (E) average EHF threshold difference between left and right ears. r = Spearman’s correlation, shaded area indicates 95 % confidence interval of the regression line.

## 4. Discussion

The present study assessed behavioral and electrophysiological assays of hearing integrity in young, normal-hearing listeners with normal audiograms (i.e., 250-8000 Hz) but who varied in their EHF thresholds (9-16 kHz). Listeners with elevated EHF thresholds were considered the “high-risk” group, suspected of degeneration in the cochlear synapses characteristic, i.e. cochlear synaptopathy^8^. We presented single- and paired-clicks to both low- and high-risk participants to test if differential changes in the cochlear-initiated SP potential with ICI would differentiate hearing groups and/or participants with degraded auditory processing as indexed by perceptual speech-in-noise measures.

Unlike most studies which measured SP by human visual inspection^7–9^, we automated our SP measurements by fitting a logarithmic curve preceding ABR wave I through the SP “pedestal”?) to estimate the SP inflection point (Fig. 2A). This method not only saves time but also reduced human error due to inter- and intra-observers’ variability. Additionally, SP amplitudes to paired-clicks were normalized to those of single-clicks to minimize the impact of inter-subject differences in head size, tissue conductivity, etc., that normally confound absolute measures of ECochG quantification. Critically, we observed an increase in SP amplitudes evoked by paired-clicks compared to standard single-clicks in high-risk listeners (Fig. 2B). However, less changes in SP amplitudes were found in low-risk listeners when stimulated with either single- or paired-clicks. SP ratios were also predictive of behavioral SIN perception (3A-B) and subjective hearing acuity (3C) confirming the behavioral relevance of our electrophysiological markers.

### 4.1 SP responses evoked by single- vs. paired-click

ABR wave I (i.e., the action potential (AP) of the ECochG) originates from the cochlear nerve. The single-fiber AP component is an “all-or-none” response which exhibits neural refractoriness to a preceding stimulus. There are two observed refractory mechanisms: the absolute refractory period and the relative refractory period. The absolute refractory period of a human cochlear nerve AP is reported to be ~1.0 ms^25^ to 1.2 ms^31^. Following AP generation to a first click, the auditory nerve cannot again discharge to a second click within this absolute refractory period. During the relative refractory period, it is difficult, but not impossible, for the cochlear nerve fibers to discharge and generate new APs as it is characterized by an exponentially decreasing threshold for the AP to a resting level with a time constant of 4-5 ms^25^. Unlike the AP (i.e., ABR wave I), the SP is a receptor response originating from the hair cells. Consequently, it is not constrained by neural refractoriness. Two types of potentials related to hair cells can contribute to SP: 1) non-linear components in the receptor potentials from inner and/or outer hair cells that are not removed by alternating stimulus polarity, and 2) EPSPs from cochlear nerve terminals under the inner hair cells^8^. It is possible that the loss of a negative EPSP could contribute to SP enhancement in listeners who have putative cochlear synaptopathy and attenuated MEMR, as reported in synaptopathic mice^22^. As the SP is not constrained by refractoriness, paired-clicks with ICIs within the absolute refractory period can thus elicit additional and enhanced SP responses^8^. While we did not find attenuated ARTs in our listeners, we suggest the former mechanism might account for the larger SPs we found in high-risk ears.

### 4.2 Differential changes in SP relate to hearing acuity and speech-in-noise perception

Our results show that SP ratios of rapid auditory stimuli (ICI = 0.1 and 0.2 ms) correlated with QuickSIN (Fig. 3A-B) while SP ratios for ICI = 1.0 ms correlated with subjective acuity rating (Fig. 3C). The rapid ICIs (0.1-0.2 ms) fall within the absolute refractory period of the auditory nerve while an ICI of 1.0 ms is closer to the limit of absolute refractory period. In addition to not being influenced by neural refractoriness, larger SP ratios for very rapid ICIs in listeners with worse SIN perception (i.e., larger SNR-loss) could be due to unusually small synaptic adaptation in terms of neurotransmitter re-uptake. On the other hand, larger SP ratios at longer ICIs (1.0 ms), correlated with worse hearing acuity. Given that longer intervals contain neural refractory effects, this association could be related to both reduced synaptic and neural effects. It is possible our SP effects are at least partially driven by inner hair cell dysfunction in basal cochlear regions, as implied by the elevated EHF thresholds in high-risk subjects. However, hair cell impairment would tend to reduce electrophysiogical responses whereas we instead see the opposite: enlarged SPs in risk adverse ears. This argues for a neural rather than pure sensory account of our data. We speculate that, in addition to EPSPs, the reduction of synaptic adaptation due to synaptic loss may also contribute to larger SP ratios to very short ICIs in cochlear synaptopathy. In contrast, SP amplitudes to short stimuli were reported to be slightly reducing, because of adaptation, in normal-hearing humans as stimulus repetition rate increased^21^. Proper adaptation is important in maintaining normal hearing function because it is a fundamental principle of sensory processing that enables sensory information to be represented adequately^32^ and remain robust to noise^33^.

High-risk subjects in Liberman et al. (2016) who had putative cochlear synaptopathy were shown to have poorer speech discrimination scores when the task was performed under conditions of degradation (e.g., noise, time-compression, reverberation). Listeners in that study were assigned to high-risk group based on subjective responses on questionnaires related to medical history of ear and hearing function, history of noise-exposure, and use of hearing protection. In contrast, in this study, we defined high-risk listeners objectively based on their average EHF thresholds and not noise-exposure history. Arguably, the stark differences in EHFs (without elevated thresholds in the normal audiometric range) suggest the hearing loss in our high-risk ears was far from “hidden.” Still, the association of SP and QuickSIN score or subjective acuity rating in this study and poorer SIN performance in high-risk group of Liberman’s study suggest that recruiting and assigning subjects into different hearing risk groups based on more difficult SIN perceptual tests or more detailed subjective responses regarding hearing and historical noise-exposure might be a better approach.

### 4.3 SP versus ABR measures of cochlear synaptopathy

Although ABR wave I amplitudes are usually reduced by neural damage, SP amplitudes remain robust in animals with both noise-induced and age-related synaptopathy^2^. The literature is, however, highly equivocal and some studies fail to find associations between ABR wave I amplitude and noise exposure in humans^34–38^. Absolute amplitude of ABR wave I is highly variable in humans when recorded at the scalp. This is also true for the SP. However, the fact that SP is a receptor potential and not neural response might render it more sensitive to synaptopathic effects since these listeners usually have intact hair cells but loss of synapses. In addition, our use of *relative* (i.e., SP ratios comparing responses across stimuli) rather than absolute amplitude measures seem to make it less susceptible to confounding sources of variances and thus sensitive to putative cochlear synaptopathy.

Recently, envelope-following responses (EFRs) recorded to rectangular sinusoidal modulation (RAM) sounds was shown to predict speech-in-noise performance in normal-listening listeners with suspected cochlear synaptopathy^39^. It was claimed that RAM-EFRs yielded greater diagnostic utility than other metrics, including click-evoked ABRs and MEMRs, in predicting cochlear neural deficits in normal-hearing listeners. EFRs, however, may still be confounded by other unknown deficits in the auditory system since the response generators are located in the rostral brainstem rather than auditory periphery^40^. As such, auditory deficits reported in EFRs might not reflect degeneration in cochlear synapses or nerve fibers, per se, but rather more central effects that are entirely independent of cochlear synaptopathy. In contrast, the source of SP is specifically localized to cochlear processes, which are the central etiology of cochlear synaptopathy. As a result, the more direct interpretation of the SP renders it a higher potential than EFRs to be used as a diagnostic tool for cochlear synaptopathy and related neural degeneration.

Lastly, sample size limitations of this study are worth mention. While the group differences and correlational effects observed here (e.g., *η_p_*^2^ > 0.22; r > 0.55-0.6) are considered intermediate to large effects^41^ and thus provide moderate to strong evidence favoring the alternative hypothesis, we acknowledge the limitation of our smaller sample size. Additional studies with larger populations of low- vs. high-risk listeners are needed to replicate and confirm the preliminary findings of our proof-of-concept measures. Still, the convergence of group differences across several behavioral and electrophysiological measures is promising.

## Acknowledgments

We would like to thank Kate R. Allen and Ashley A. Peeples for assistance with data collection. This work was supported by the National Institutes of Health (NIH/NIDCD R01DC016267) (G.M.B.).

